# Genomic diversity of *Escherichia coli* isolates from backyard chickens and guinea fowl in the Gambia

**DOI:** 10.1101/2020.05.14.096289

**Authors:** Ebenezer Foster-Nyarko, Nabil-Fareed Alikhan, Anuradha Ravi, Nicholas M. Thomson, Sheikh Jarju, Brenda Anna Kwambana-Adams, Arss Secka, Justin O’Grady, Martin Antonio, Mark J. Pallen

## Abstract

Chickens and guinea fowl are commonly reared in Gambian homes as affordable sources of protein. Using standard microbiological techniques, we obtained 68 caecal isolates of *Escherichia coli* from ten chickens and nine guinea fowl in rural Gambia. After Illumina whole-genome sequencing, 28 sequence types were detected in the isolates (four of them novel), of which ST155 was the most common (22/68, 32%). These strains span four of the eight main phylogroups of *E. coli*, with phylogroups B1 and A being most prevalent. Nearly a third of the isolates harboured at least one antimicrobial resistance gene, while most of the ST155 isolates (14/22, 64%) encoded resistance to ≥3 classes of clinically relevant antibiotics, as well as putative virulence factors, suggesting pathogenic potential in humans. Furthermore, hierarchical clustering revealed that several Gambian poultry strains were closely related to isolates from humans. Although the ST155 lineage is common in poultry from Africa and South America, the Gambian ST155 isolates belong to a unique cgMLST cluster comprised of closely related (38-39 alleles differences) isolates from poultry and livestock from sub-Saharan Africa—suggesting that strains can be exchanged between poultry and livestock in this setting. Continued surveillance of *E. coli* and other potential pathogens in rural backyard poultry from sub-Saharan Africa is warranted.

**Author notes:** All supporting data and protocols have been provided within the article or as supplementary data files. Eleven supplementary figures and eight supplementary files are available with the online version of this article.

**Data summary:** The genomic assemblies for the isolates reported here are available for download from EnteroBase (http://enterobase.warwick.ac.uk/species/index/ecoli) and the EnteroBase assembly barcodes are provided in File S2.

Sequences have been deposited in the NCBI SRA, under the BioProject ID: PRJNA616250 and accession numbers SAMN14485281 to SAMN14485348 (File S2). Assemblies have been deposited in GenBank under the BioProject ID: PRJNA616250 and accession numbers CP053258 and CP053259.

**Impact statement:** Domestic birds play a crucial role in human society, in particular contributing to food security in low-income countries. Many households in Sub-Saharan Africa rear free-range chickens and guinea fowl, which are often left to scavenge for feed in and around the family compound, where they are frequently exposed to humans, other animals and the environment. Such proximity between backyard poultry and humans is likely to facilitate transmission of pathogens such as *Escherichia coli* or antimicrobial resistance between the two host species. Little is known about the population structure of *E. coli* in rural chickens and guinea fowl, although this information is needed to contextualise the potential risks of transmission of bacterial strains between humans and rural backyard poultry. Thus, we sought to investigate the genomic diversity of *E. coli* in backyard poultry from rural Gambia.

## Introduction

The domestic chicken (*Gallus gallus domesticus)* is the most numerous bird on the planet, with an estimated population of over 22.7 billion—ten times more than any other bird [1]. Since their domestication from the red jungle fowl in Asia between 6, 000 and 8, 000 years ago [2, 3], chickens have been found almost everywhere humans live. Other poultry, such as turkeys, guinea fowl, pheasants, duck and geese, are derived from subsequent domestication events across Africa, Europe and the Americas [4]. For example, the helmeted guinea fowl (*Numida meleagris)* originated in West Africa, although domesticated forms of this bird are now found in many parts of the tropics.

Poultry are reared for meat, eggs and feathers [5]. Poultry production is classified into four sectors, based on the marketing of poultry products and the level of biosecurity [6]. Intensive poultry farming falls under sectors 1 to 3, characterised by moderate to high levels of biosecurity, while sector 4 pertains to the “backyard”, “village” or “family” poultry system, with little or no biosecurity measures.

In rural backyard farming—prevalent in low- to middle-income countries such as the Gambia—a small flock of birds (between one and fifty) usually from indigenous breeds are allowed to scavenge for feed over a wide area during the daytime, with minimal supplementation, occasional provision of water and natural hatching of chicks. The poultry may be confined at night in rudimentary shelters to minimise predation, or birds may roost in owners’ kitchens, family dwellings, tree-tops, or nest in the bush [7]. Urban and peri-urban backyard poultry farming—for example, in Australia, New Zealand (North Island), the US and in the UK—differs from rural backyard farming in that the birds are kept on an enclosed residential lot [8-10].

Backyard poultry fulfils important social, economic and cultural roles in many societies. Seventy percent of poultry production in low-income countries comes from backyard poultry [11]. The sale of birds and eggs generates income, while occasional consumption of poultry meat provides a source of protein in the diet. In traditional societies, domestic poultry meat is considered tastier than commercial broiler meat and, as it is perceived to be tougher in texture, is preferred for preparing dishes that require prolonged cooking [12]. It is estimated that meat and eggs from backyard poultry contribute about 30% of the total animal protein supply of households in low-income countries [13, 14]. In rural Gambia, backyard poultry can be offered as gifts for newlyweds or sold to solve family needs such as paying school fees, buying new clothes or other household needs [7]. Chickens may also be used as offering to a traditional healer, consumed when there is a guest, or during ceremonies. Urban and peri-urban poultry are kept mostly for home consumption of their eggs or meat, but also as pets or used for pest control [8, 15-17]. The proximity between backyard poultry and humans may facilitate transmission of pathogens such as *Escherichia coli* between the two host species.

*E. coli* is a generalist bacterium that commonly colonises the gastrointestinal tract of mammals and avian species [18]. Based on their pathogenic potential, *E. coli* can be divided into three categories: commensals, diarrhoeagenic *E. coli* and extraintestinal pathogenic *E. coli* (ExPEC). ExPEC frequently colonise the gut asymptomatically; however, they possess a wide range of unique virulence factors that enable them to colonise extraintestinal tissues in humans, pets and poultry [19, 20]. A sub-pathotype of ExPEC strains, known as Avian Pathogenic *E. coli* (APEC), cause colibacillosis—an extraintestinal disease in birds, with manifestations such as septicaemia, air sacculitis and cellulitis [21]. Avian colibacillosis results in high mortality and condemnation of birds, resulting in significant annual economic losses for the poultry industry [22]. As a result, antimicrobials are often used in intensive farming systems to prevent bacterial infections and treat sick birds—a practice that has been linked to the development of antimicrobial resistance (AMR) in poultry.

Although previous studies have focused on the detection of AMR and documented the emergence of multiple-drug resistance (MDR) in this niche [23-27], little is known about the population structure of *E. coli* in rural backyard poultry. Given the increased exposure to humans, the natural environment and other animals, the population of *E. coli* in birds raised under the backyard system may differ considerably from those reared in intensive systems. It is also possible that the lineages of *E. coli* within local genotypes of rural poultry might differ between geographical regions. The absence of biosecurity measures in backyard poultry farming increases the potential for zoonotic transmission of pathogenic and/or antimicrobial-resistant strains to humans.

In a recent study of commercial broiler chickens, multiple colony sampling revealed that a single broiler chicken could harbour up to nine sequence types of *E. coli* [28]. However, within-host diversity of *E. coli* in backyard poultry—particularly in guinea fowl—has not been well studied and so we do not know how many lineages of *E. coli* can co-colonise a single backyard bird. To address these gaps in our knowledge, we exploited whole-genome sequencing to investigate the genomic diversity and burden of AMR among *E. coli* isolates from backyard chickens and guinea fowl in rural Gambia, West Africa.

## Materials and methods

### Study population

The study population comprised ten local-breed chickens and nine guinea fowl from a village in Sibanor in the Western Region of the Gambia (Table 1). Sibanor covers an area of approximately 90 km^2^ and is representative of rural areas in the Gambia [29]. It has a population of about 10,000. Most of the villagers are subsistence farmers growing peanuts, maize and millet. Households within this community comprise extended family units of up to fifteen people, which make up the “compound”. All guinea fowl were of the pearl variety, characterised by purplish-grey feathers dotted with white.

**Table 1:**
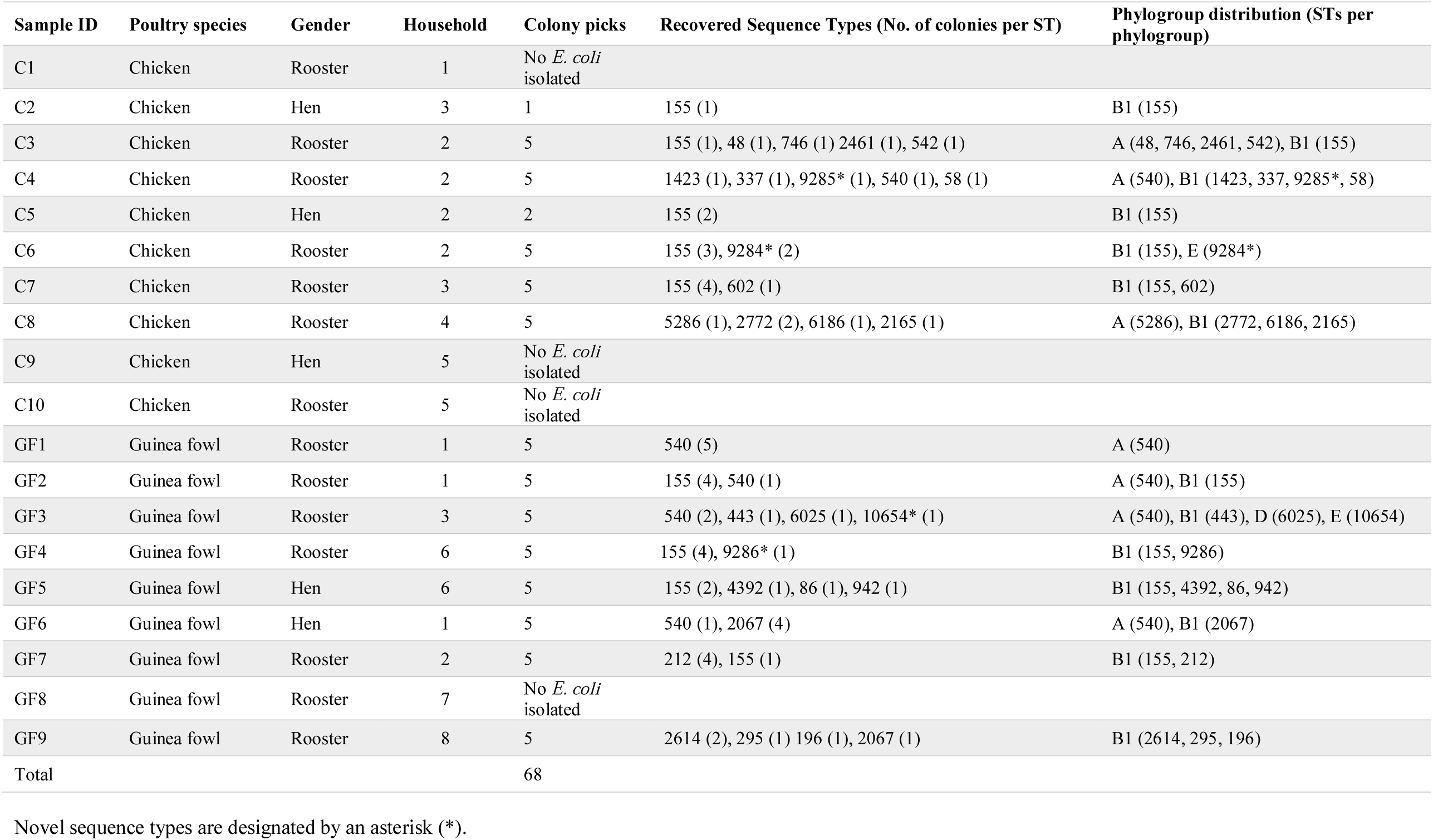
Characteristics of the study population

### Sample collection

The sampling was done in November 2016. Poultry birds were first observed in motion for the presence of any abnormalities. Healthy-looking birds were procured from eight contiguous households within 0.3-0.4 km of each other and transported to the Abuko Veterinary Station, the Gambia in an air-conditioned vehicle. A qualified veterinarian then euthanised the birds and removed their caeca under aseptic conditions. These were placed into sterile falcon tubes and flash-frozen on dry ice in a cooler box. The samples were transported to the Medical Research Council Unit The Gambia at the London School of Hygiene and Tropical Medicine labs in Fajara, where the caecal contents were aseptically emptied into new falcon tubes for storage at -80°C within 3 h. A peanut-sized aliquot was taken from each sample into a 1.8 ml Nunc tube containing 1 ml of Skimmed Milk Tryptone Glucose Glycerol (STGG) transport and storage medium (Oxoid, Basingstoke, UK), vortexed at 4200rpm for 2 min and frozen at -80°C. Figure 1 summarises the sample processing flow.

**Figure 1.**
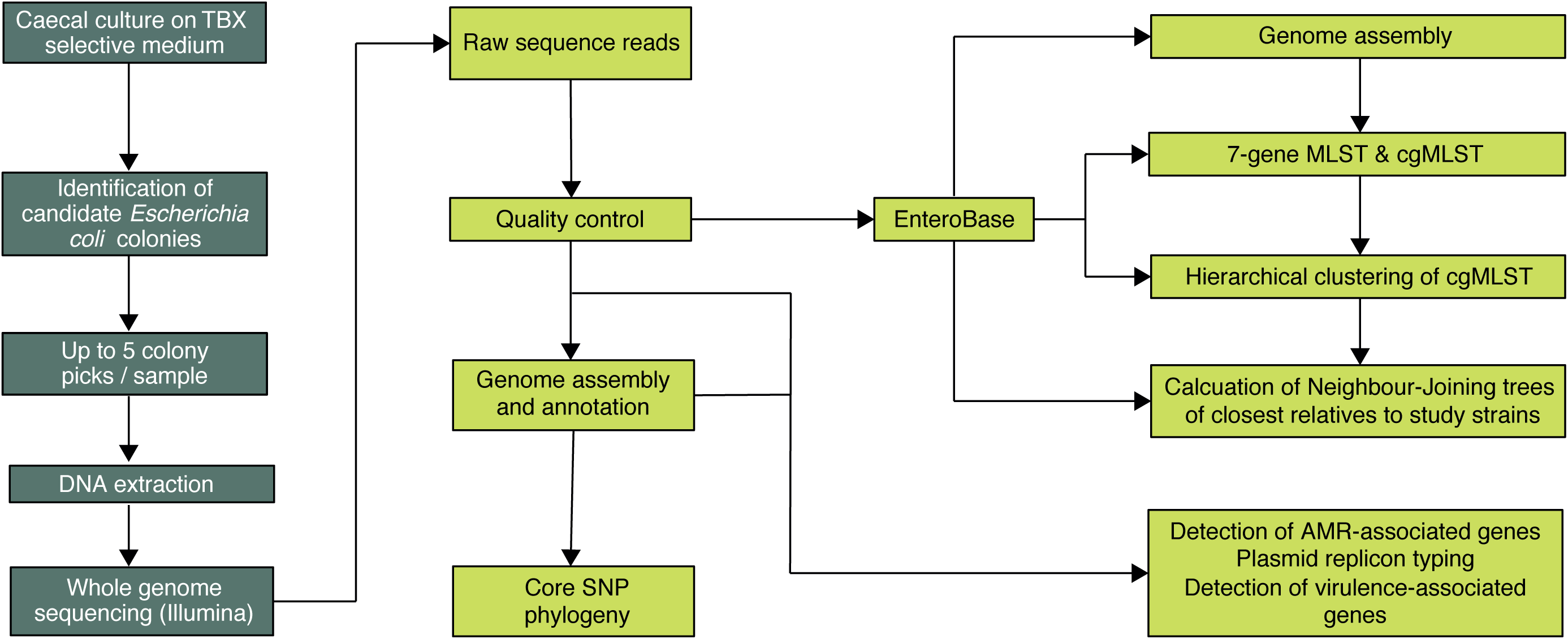
Study sample-processing flow diagram. TBX, Tryptone-Bile-X-glucoronide agar; MLST, multi-locus sequence typing; cgMLST, core genome multi-locus sequence typing.

### Microbiological processing

Isolation of *E. coli* was performed as follows. The caecal-STGG suspension was removed from -80 °C storage and allowed to thaw briefly on wet ice. A 100 µl aliquot was then taken into 900 µl of physiological saline (0.85%) and taken through four ten-fold serial dilutions. A100 µl aliquot each was then taken from the dilutions and uniformly streaked onto tryptone-bile-X-glucoronide agar plates using the spread plate technique. The inoculated plates were incubated at 37°C for 18–24 h under aerobic conditions. Following overnight incubation, colony counts were determined for raised, translucent and entire colonies that exhibited bluish-green pigmentation typical of *E. coli*. Up to five candidate colonies were selected per sample and sub-cultured on MacConkey agar. These were incubated at 37°C in air for 18–24 h and stored in 20% glycerol broth at -80°C. The isolates from chickens were designated “C1-C10”, while those from guinea fowl were prefixed by “G1-G9”, followed by the respective colony number (1 up to 5).

### Genomic DNA extraction

Genomic DNA was extracted from overnight broth cultures prepared from each single colony sub-culture using the 96-well plate lysate method as described previously [30]. The DNA was eluted in Tris-Cl (pH, 8.0) and quantified using the Qubit high sensitivity DNA assay kit (Invitrogen, MA, USA). DNA samples were kept at -20°C until Illumina sequencing library preparation. Broth cultures were spun at 3500rpm for 2 min and lysed using lysozyme, proteinase K, 10% SDS and RNase A in Tris EDTA buffer (pH 8.0).

### Illumina sequencing

Whole-genome shotgun sequencing of the DNA extracts was performed on the Illumina NextSeq 500 instrument (Illumina, San Diego, CA) using a modified Illumina Nextera library preparation protocol as described previously [30]. We run the final pooled library at a concentration of 1.8 pM on a mid-output flow cell (NSQ® 500 Mid Output KT v2 300 cycles; Illumina Catalogue No. FC-404-2003) according to manufacturer’s instructions. Following sequencing, FASTQ files were downloaded from BaseSpace to a local server hosted at the Quadram Institute Bioscience.

### Genome assembly and phylogenetic analysis

The raw sequences were initially analysed on the Cloud Infrastructure for Microbial Bioinformatics [31]. This included concatenating paired-end short reads, quality checks with FastQC v0.11.7 [32], trimming with Trimmomatic v0.39 [33] and assembly by Spades v3.13.2 [34]. The quality of the assemblies was checked using QUAST v5.0.0, de6973bb [35] and annotation of the draft genomes was carried out using Prokka v1.13.3 [36]. We used the mlst software (https://github.com/tseemann/mlst) to call multi-locus sequence types (MLSTs) using the Achtman scheme [37], based on the seven house-keeping genes, *adk, fum*C, *gyr*B, *icd, mdh, pur*A and *rec*A. We used Snippy v4.3.2 (https://github.com/tseemann/snippy) for variant calling and to generate a core-genome alignment, from which a maximum-likelihood phylogenetic tree was reconstructed using RAxML v8.2.4 [38], based on a general time-reversible nucleotide substitution model with 1,000 bootstrap replicates. We included representative reference genome sequences for the major phylogroups of *E. coli* and *Escherichia fergusonii* as an outgroup (File S1). Given that recombination is widespread in *E. coli* and tends to blur phylogenetic signals [37], we used Gubbins (Genealogies Unbiased By recomBinations In Nucleotide Sequences) [39] to detect and mask recombinant regions of the core-genome alignment prior to the phylogenetic reconstruction. We used the GrapeTree [40] to visualise and annotate phylogenetic trees. We calculated pair-wise single nucleotide polymorphism (SNP) distances between genomes from the core-genome alignment using snp-dists v0.6 (https://github.com/tseemann/snp-dists).

Subsequently, the short-read sequences were uploaded to EnteroBase [41], an online genome database and integrated software environment which currently hosts more than 138,164 *E. coli* genomes, sourced from all publicly available sequence databases and user uploads. EnteroBase routinely retrieves short-read *E. coli* sequences from the public domain, performs quality control and *de novo* assemblies of Illumina short-read sequences, annotates these and assigns seven-allele MLST (ST) and phylogroups from genome assemblies using standardised pipelines. In addition, EnteroBase assigns unique core-genome MLST (cgMLST) numbers to each genome, based on the typing of 2,512 genes in *E. coli.*

### Population structure analysis

We utilised the Hierarchical Clustering (HierCC) algorithm in EnteroBase to assign our poultry genomes to stable clusters designated as HC0 up to HC1100, based on pair-wise differences between genomes at cgMLST alleles. In *E. coli*, the HC400 cluster has been shown to correspond to strain endemicity, while HC1100 corresponds to the seven-allele MLST clonal complexes [41]. The HierCC algorithm therefore lends itself as a very useful tool for the analysis of bacterial population structures at multiple levels of resolution. In a recent study of the population structure of *Clostridioides difficile*, Frentrup et al [42] showed that HierCC allows closely-related neighbours to be detected at 89% consistency between cgMLST pair-wise allelic differences and SNPs. We determined the closest relatives to our study *E. coli* isolates using the HC1100 cluster and reconstructed neighbour-joining trees using NINJA [43]. In order to compare the strain distribution that we observed among our study isolates with what pertains in poultry *E. coli* isolates from elsewhere, we further retrieved genomic assemblies from all publicly-available poultry *E. coli* isolates, stratified by their source continent and reconstructed NINJA neighbour-joining trees depicting the prevalence of STs per continent.

### Analysis of accessory gene content

We used ARIBA v2.12.1 [44] to detect virulence factors, antimicrobial resistance genes and plasmid replicons. Briefly, this tool scans the short-read sequences against the core Virulence Factors Database [45] (virulence factors), ResFinder (AMR) [46] and PlasmidFinder (plasmid-associated genes) [47] databases and generates customised outputs, based on a percentage identity of ≥ 90% and coverage of ≥ 70%. The VFDB-core, ResFinder and PlasmidFinder databases were downloaded on 29 October 2018. Virulence factors were visualised by overlaying them onto the phylogenetic tree using the ggtree, ggplot2 and phangorn packages in RStudio v3.5.1.

We determined the prevalence of AMR genes among poultry *E. coli* isolates from the rest of the world, for comparison with what we found in isolates from this study. To do this, we interrogated the downloaded continent-stratified genomes as above using ABRicate v0.9.8 (https://github.com/tseemann/abricate) to predict AMR-associated genes by scanning against the ResFinder database (accessed 28 July 2019), based on a percentage identity threshold of ≥ 90% and a coverage of ≥ 70%.

### Antimicrobial susceptibility

Due to logistic constraints, a third of the study isolates (20/68, 29%) were randomly selected for phenotypic susceptibility testing by minimum inhibitory concentrations (MICs). MICs were performed by the agar dilution method [48], according to the European Committee on Antimicrobial Susceptibility Testing v. 9.0 (EUCAST, 2019) guidelines. Stock solutions of 1,000 mg l^-1^ were initially prepared, from which the working solutions were made. For each antibiotic, duplicate two-fold serial dilutions (from 32mg/L to 0.03 mg l^-1^) were done in molten Mueller-Hinton agar (Oxoid, Basingstoke, UK). The results were interpreted according to EUCAST breakpoint tables (http://www.eucast.org). Where EUCAST cut-off values were not available, the recommended cut-off values from the Clinical Laboratory Standards Institute (https://www.clsi.org) (Performance Standards for Antimicrobial Susceptibility Testing (28^th^ Information Supplement, M100-S28) were used.

### Oxford nanopore sequencing

Prior to sequencing, DNA fragments were assessed using the Agilent 2200 TapeStation (Agilent Catalogue No. 5067-5579) to determine the fragment lengths. Long-read sequencing was carried out using the rapid barcoding kit (Oxford Nanopore Catalogue No. SQK-RBK004). Libraries were prepared following the manufacturer’s instructions. An input DNA concentration of 400 ng was used for the library preparation and a final concentration of 75 µl of the prepared library loaded onto an R9.4 MinION flow cell. The final concentration of the library pool was assessed using the Qubit high-sensitivity DNA assay (Invitrogen, MA, USA).

### Hybrid assembly and analysis of plasmids and phages

The long reads were base-called with Guppy, the Oxford Nanopore Technologies’ post-sequencing processing software (https://nanoporetech.com/). The base-called FASTQ files were then concatenated into a single file each and demultiplexed based on their respective barcodes, using the qcat python command-line tool v1.1.0 (https://github.com/nanoporetech/qcat). We performed hybrid assemblies of the Illumina and nanopore reads with Unicycler v0.4.8.0 [49]. The quality of the hybrid assemblies was assessed with QUAST v5.0.0, de6973bb [35]. The hybrid assemblies were analysed for the presence of plasmids and prophages using ABRicate PlasmidFinder and PHASTER [50] respectively. Annotations of the assemblies were carried out using Prokka v1.13.3 [36].

### Ethical statement

The study was approved by the joint Medical Research Council Unit The Gambia-Gambian Government ethical review board.

## Results

### Study population

We analysed nineteen caecal samples obtained from ten chickens and nine guinea fowl. Fifteen out of the nineteen (79%) samples yielded growth of *E. coli* on culture, from which 68 colonies were recovered.

### Sequence type and phylogroup distribution

We recovered 28 seven-allele sequence types (STs), of which ST155 was the most common (22/68, 32%). Four of the STs were novel—two from chickens and two from guinea fowl. Seventeen of the 28 STs have been previously isolated from humans and or other vertebrates, while five (ST942, ST2165, ST2461, ST4392 and ST5826) have not been seen in humans before (Table 2). The isolates were spread over four of the eight known phylogroups of *E. coli*, but most belonged to phylogroups B1 and A, which are home to strains associated with human intestinal infections and avian colibacillosis [51, 52] (Figure 2). Hierarchical clustering resolved the study strains into 22 cgMLST complexes, indicating a high level of genomic diversity (File S2).

**Table 2:** Prevalence of the study sequence types in EnteroBase

| ST | Source | Phylotype | Prevalence in EnteroBase |
| --- | --- | --- | --- |
| 48 | Chicken | A | Human, livestock, *Celebes* ape |
| 58 | Chicken | B1 | Human, livestock, poultry |
| 86 | Guinea fowl | B1 | Human, livestock, companion animal, poultry |
| 155 | Chicken, Guinea fowl | B1 | Human, poultry, mink, livestock |
| 196 | Guinea fowl | B1 | Human, livestock, companion animal, environment |
| 212 | Guinea fowl | B1 | Human, poultry, deer, companion animal |
| 295 | Guinea fowl | B1 | Human, poultry, livestock, companion animal, environment, food, |
| 337 | Chicken | B1 | Human, rhinoceros, poultry, environment (soil and water) |
| 443 | Guinea fowl | B1 | Human, environment, livestock |
| 540 | Chicken, Guinea fowl | A | Human, environment (water and sewage), livestock, poultry, gull, rabbit, plant, oyster, fish |
| 542 | Chicken | A | Human, livestock, poultry |
| 602 | Chicken | B1 | Human, poultry, livestock, bird, fish, reptile |
| 746 | Chicken | A | Human, poultry, fish, livestock, environment (water) |
| 942 | Guinea fowl | B1 | Environment, food, companion animal, livestock |
| 1423 | Chicken | B1 | Human, reptile, livestock |
| 2067 | Guinea fowl | B1 | Human, environment |
| 2165 | Chicken | B1 | Livestock, companion animal, reptile, bird |
| 2461 | Chicken | A | Sheep, poultry |
| 2614 | Guinea fowl | B1 | Human |
| 2772 | Chicken | B1 | Human, livestock, environment |
| 4392 | Guinea fowl | B1 | Livestock, wild animal, companion animal |
| 5826 | Chicken | A | Poultry |
| 6025 | Guinea fowl | D | Unknown source§ |
| 6186 | Chicken | B1 | Livestock, environment |
| 9284 | Chicken | E | Novel |
| 9285 | Chicken | B1 | Novel |
| 9286 | Guinea fowl | B1 | Novel |
| 10654 | Guinea fowl | D | Novel |

§ ST6025 occurred only one other isolate in EnteroBase, beside the study ST6025 strain. However, the source of isolation of this other isolate was not available.

**Figure 2.**
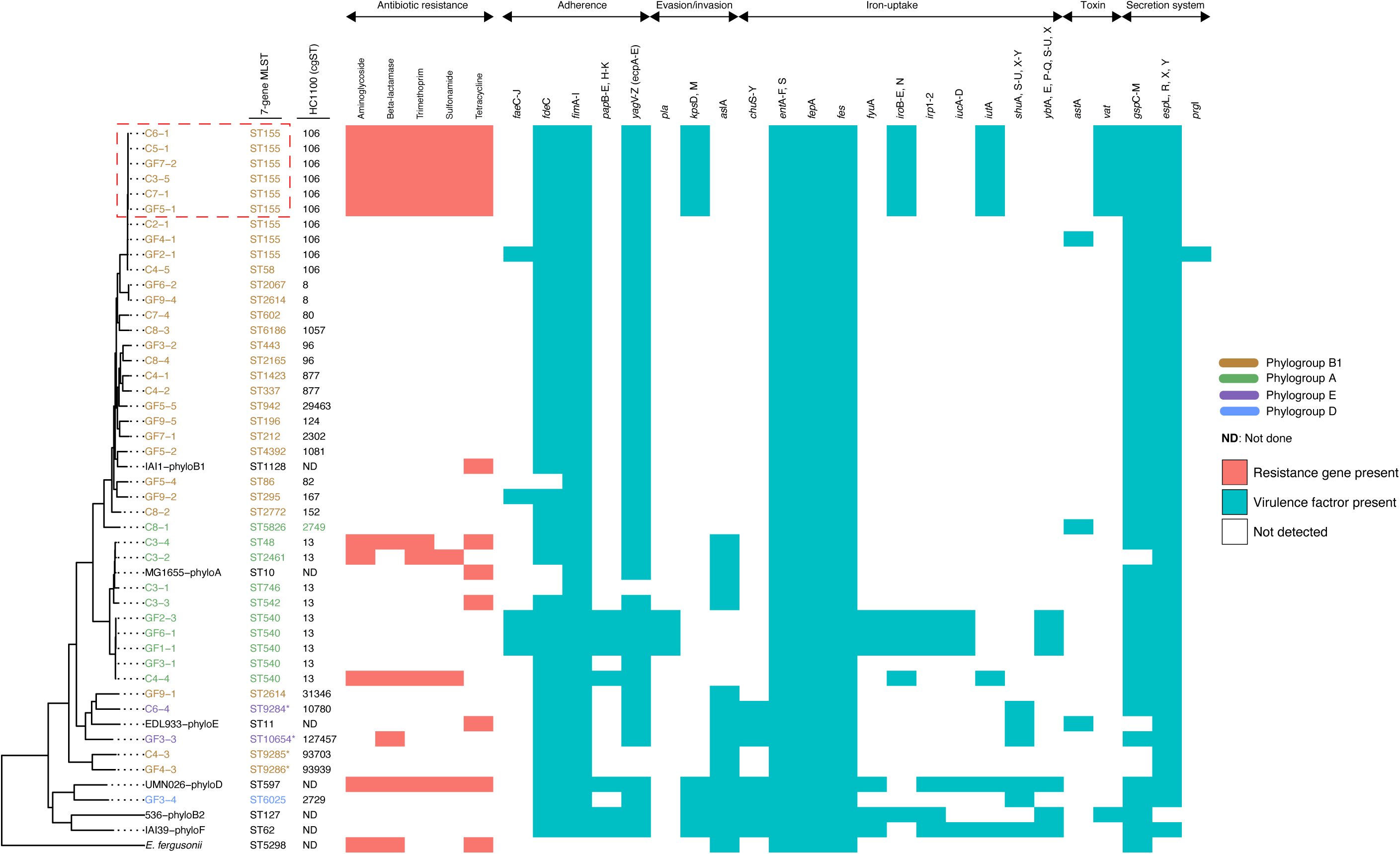
A maximum-likelihood phylogeny of the study isolates reconstructed with RAxML, based on non-repetitive, non-recombinant core SNPs, using a general time-reversible nucleotide substitution model with 1,000 bootstrap replicates. The tip labels indicate the sample names, with the respective Achtman sequence types (ST) and HC1100 (cgST complexes) indicated next to the sample names. The colour codes indicate the respective phylogroups to which the isolates belong. The outgroup and the other *E. coli* reference genomes denoting the major *E. coli* phylogroups are in black. Asterisks (*) are used to indicate novel STs. Overlaid on the tree are the predicted virulence factors for each isolate. The virulence genes are grouped according to their function. Chicken isolates are denoted “C” and guinea fowl samples by “GF”, with the suffix indicating the colony pick. We have not shown multiple colonies of the same Achtman ST recovered from a single individual—in such instances, only one representative isolate is shown. Nor have we shown virulence factors that were detected only in the reference genomes. The red box highlights multi-drug resistant isolates which concurrently harbour putative fitness and colonisation factors that are important for invasion of host tissues and evasion of host immune defences. The full names of virulence factors and their known functions are provided in File S8.

We generated complete, circular genome assemblies of the two novel sequence types isolated from guinea fowl: ST10654 (GF3-3) and ST9286 (GF4-3). Although neither strain encoded AMR genes or plasmids, GF3-3 contained three prophages (two intact, one incomplete), while GF4-3 harboured four prophages (three intact, one incomplete) (File S3).

### Within-host genomic diversity and transmission of strains

Several birds (12/19, 63%) were colonised by two or more STs; in most cases, the STs spanned more than two phylotypes (Table 1). In two chickens, all five colony picks belonged to distinct STs. We observed some genetic diversity among multiple colonies of the same ST recovered from the same host (Table 3A). Most of these involved variants that differed by 0-4 SNPs, i.e. variation likely to have arisen due to within-host evolution. However, in one instance, pair-wise SNP differences (ranging from 4 to 255) suggested independent acquisition of distinct clones. Pair-wise SNP analysis also suggested transmission of strains—including MDR isolates—between chickens and between chicken and guinea fowl (Table 3B and 3C) from the same household (File S4).

**Table 3A:**
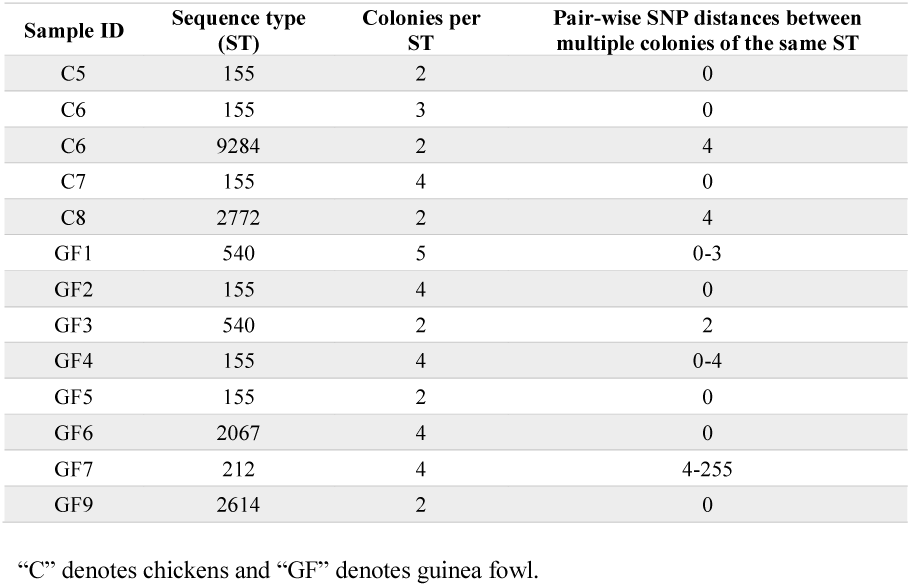
Within-host single nucleotide polymorphism diversity between multiple genomes of the same ST recovered from the same bird

**Table 3B:** Single nucleotide polymorphism differences between isolates recovered from Chicken 3, Chicken 5, Chicken 6 and Guinea fowl 7

|  | C3-5 | C5-1 | C5-2 | GF7-2 | C6-1 | C6-2 | C6-3 |
| --- | --- | --- | --- | --- | --- | --- | --- |
| C3-5 | 0 | 0 | 0 | 0 | 0 | 0 | 0 |
| C5-1 | 0 | 0 | 0 | 0 | 0 | 0 | 0 |
| C5-2 | 0 | 0 | 0 | 0 | 0 | 0 | 0 |
| GF7-2 | 0 | 0 | 0 | 0 | 0 | 0 | 0 |
| C6-1 | 0 | 0 | 0 | 0 | 0 | 0 | 0 |
| C6-2 | 0 | 0 | 0 | 0 | 0 | 0 | 0 |
| C6-3 | 0 | 0 | 0 | 0 | 0 | 0 | 0 |

Chicken samples are denoted by “C” and guinea fowl samples by “GF”. All the isolates in this transmission network encoded resistance to $\geq 3$ classes of antimicrobials.

**Table 3C:** Single nucleotide diversity differences between isolates recovered from guinea fowls 1, 2 and 6.

|  | GF1-1 | GF1-2 | GF1-3 | GF1-4 | GF1-5 | GF2-3 | GF6-1 |
| --- | --- | --- | --- | --- | --- | --- | --- |
| GF1-1 | 0 | 2 | 3 | 1 | 1 | 2 | 3 |
| GF1-2 | 2 | 0 | 3 | 1 | 1 | 2 | 3 |
| GF1-3 | 3 | 3 | 0 | 2 | 2 | 3 | 2 |
| GF1-4 | 1 | 1 | 2 | 0 | 0 | 1 | 2 |
| GF1-5 | 1 | 1 | 2 | 0 | 0 | 1 | 2 |
| GF2-3 | 2 | 2 | 3 | 1 | 1 | 0 | 3 |
| GF6-1 | 3 | 3 | 2 | 2 | 2 | 3 | 0 |

### Prevalence of AMR, virulence factors and plasmid replicons among the study isolates

Twenty isolates (20/68, 29%) harboured at least one AMR gene and sixteen (16/68, 24%) were MDR, i.e. positive for genes predicted to convey resistance to three or more classes of antibiotics (Figure 4; File S5). Fourteen of the sixteen MDR isolates belonged to ST155— representing 64% (14/22) of the ST155 isolates recovered in this study. Notable among the resistance genes detected was the class A broad-spectrum beta-lactamase resistance (18/68, 26%). Phenotypic resistance was confirmed in >50% of the isolates tested, with an MDR rate of 75% (15/20).

**Figure 3.**
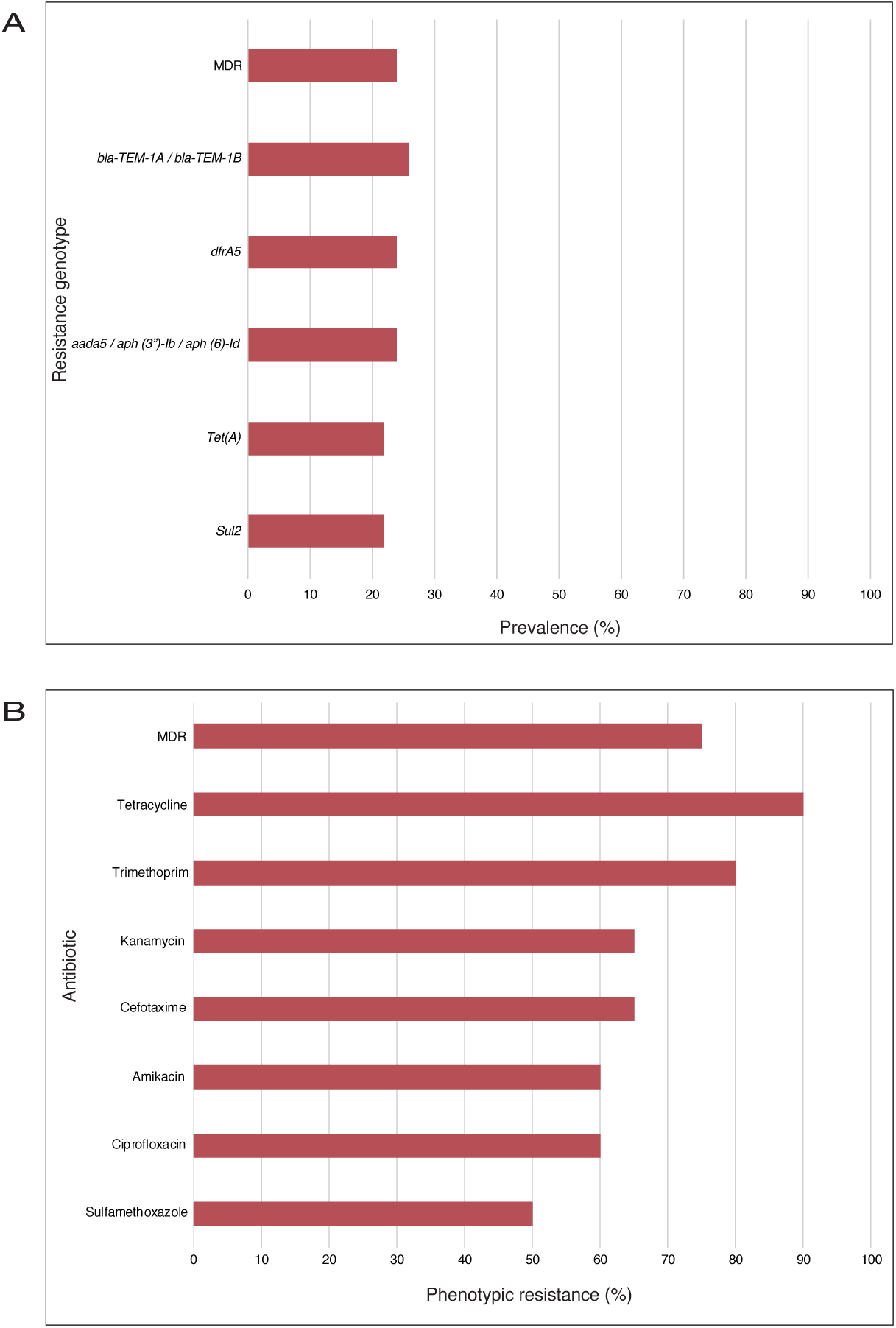
**A:** A bar graph showing the prevalence of resistance genes found among the study isolates, using the core Virulence Factors Database (reference 45) (virulence factors), ResFinder (AMR) (reference 46) and PlasmidFinder (plasmid-associated genes) (reference 47) databases, with a cut-off percentage identity of ≥ 90% and coverage of ≥ 70%. The full list of the resistance genes that were detected is presented in File S5. **B:** A bar graph depicting the prevalence of phenotypic antimicrobial resistance in 20 isolates. The results were interpreted using the recommended breakpoint tables from EUCAST (http://www.eucast.org) or the Clinical Laboratory Standards Institute (https://www.clsi.org) (Performance Standards for Antimicrobial Susceptibility Testing (28^th^ Information Supplement, M100-S28) where EUCAST cut-off values were not available.

**Figure 4.**
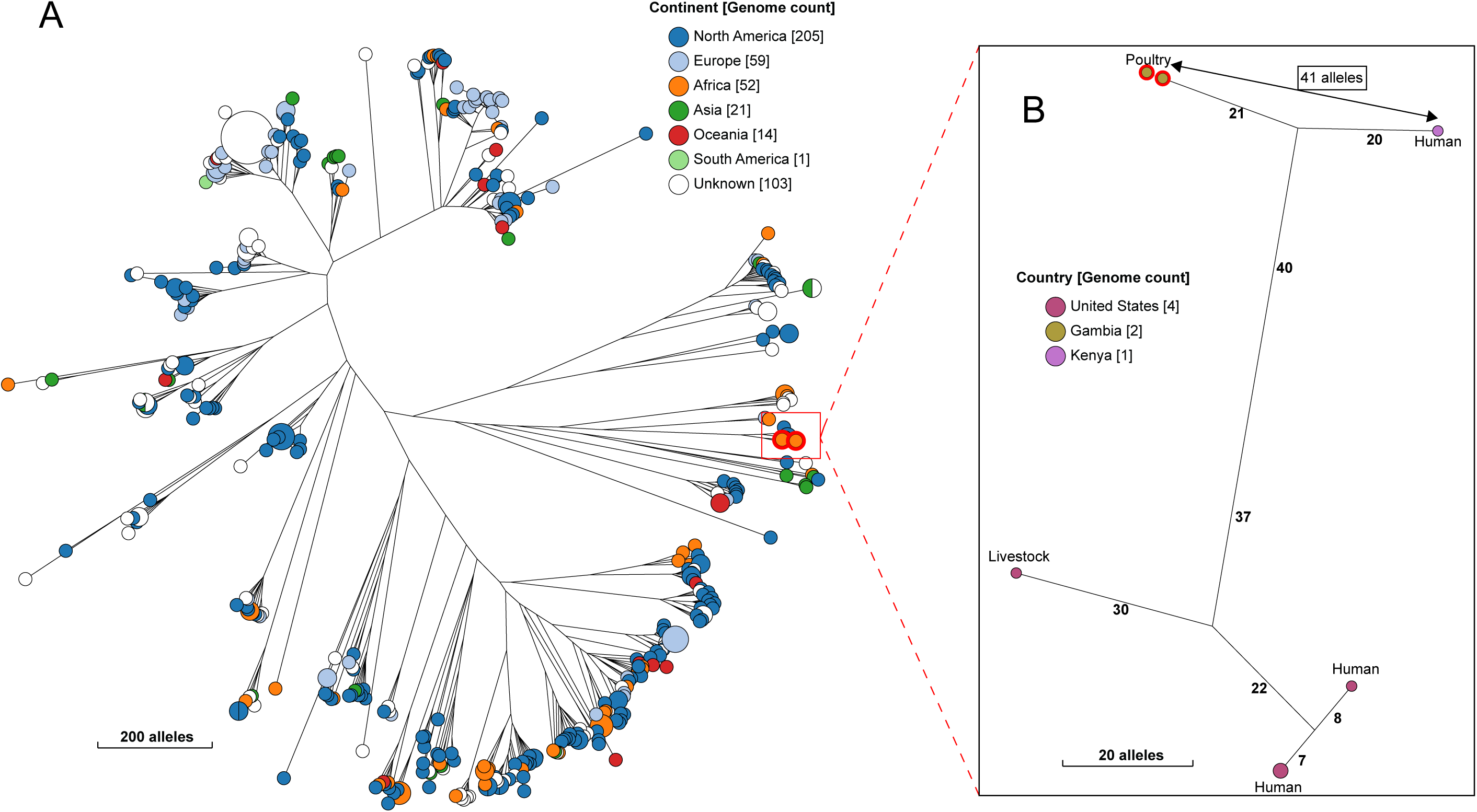
A NINJA neighbour-joining tree showing the phylogenetic relationship between our study ST2772 (Achtman) strain and all other publicly available genomes that fell within the same HC1100 cluster (cgST complex). The locations of the isolates are displayed, with the genome counts displayed in square brackets. The branch lengths are annotated with the allelic distances separating the genomes. Strains from this study are highlighted in red. The sub-tree (B) shows the closest relatives to the study strains, with the allelic distance separating them displayed with the arrow (41 alleles).

Interestingly, the MDR isolates also harboured more genes encoding putative virulence factors than did less-resistant isolates (Figure 2). Overall, 125 unique virulence-associated genes were detected from the study isolates (File S6). Notably, the virulence and AMR profiles of co-colonising STs tended to differ from each other.

One or more plasmid replicons were detected in 69% (47/68) of the study isolates—with seventeen plasmid types detected overall (File S7). IncF plasmids were the most common. A single isolate carried the col156 virulence plasmid. The multi-drug resistant isolates often co-carried large IncF plasmids (IncFIA_1, ∼27kb; IncFIB(AP001918)_1, ∼60kb; IncFIC(FII)_1, ∼56kb). Scrutiny of annotated assemblies revealed that resistance genes were often co-located on the same contig as one of the IncF plasmids. In three birds (Guinea fowl 2, Guinea fowl 5 and Guinea fowl 7), co-colonising strains (belonging to different STs) shared the same plasmid profile.

### Population dynamics of study strains

Hierarchical clustering analyses provided evidence of genomic relationships between strains from poultry and those from humans (Table 5). Significant among these were ST2772 and ST4392, which were separated from human isolates belonging to these STs by just 41 and 68 alleles in the core-genome MLST scheme respectively (Figures 4 and 5). Similarly, ST86, ST6186 and ST602 were closest to isolates from livestock (Figures S9-11), suggesting exchange of strains between livestock species.

**Table 4:** Global prevalence of AMR genes

|  | **Europe** | **Africa** | **South America** | **North America** | **Asia** | **Oceania** |
| --- | --- | --- | --- | --- | --- | --- |
| Tetracycline | 564/752, 75% | 559/591, 95% | 108/131, 83% | 2480/2975, 83% | 228/249, 92% | 132/148, 90% |
| Aminoglycoside | 303/752, 40% | 378/591, 64% | 94/131, 72% | 1497/2975, 50% | 172/249, 69% | 56/148, 38% |
| Beta-lactamase | 303/752, 40% | 246/591, 42% | 127/131, 98% | 933/2975, 31% | 157/249, 63% | 61/148, 41% |
| Sulphonamide | 338/752, 45% | 377/591, 64% | 84/131, 65% | 1174/2975, 39% | 167/249, 67% | 52/148, 35% |
| Trimethoprim | 192/752, 25% | 353/591, 52% | 58/131, 45% | 176/2975, 6% | 143/249, 57% | 66/148, 45% |
| Chloramphenicol | 303/752, 40% | 69/591, 13% | 36/131, 28% | 69/2975, 2% | 131/249, 53% | 0/148, 0% |
| Quinolone | 51/752, 7% | 144/591, 24% | 24/131, 18% | 17/2975, 1% | 74/249, 30% | 0/148, 0% |
| Lincosamide | 57/752, 8% | 0/591, 0% | 12/131, 9% | 0/2975, 0% | 14/249, 6% | 1/148, 1% |
| Macrolide | 20/752, 3% | 79/591, 13% | 3/131, 2% | 30/2975, 1% | 92/249, 37% | 0/148, 0% |
| Fosfomycin | 8/752, 1% | 4/591, 1% | 31/131, 24% | 19/2975, 1% | 71/249, 29% | 0/148, 0% |
| Streptogramin | 0/752, 0% | 0/591, 0% | 23/131, 18% | 0/2975, 0% | 0/249, 0% | 0/148, 0% |
| Colistin | 29/752, 4% | 0/591, 0% | 9/131, 7% | 0/2975, 0% | 119/249, 48% | 0/148, 0% |
| MDR | 406/752, 54% | 392/591, 66% | 100/131, 77% | 1236/2975, 42% | 175/249, 70% | 56/148, 44% |

The full list of resistance genes that were detected is presented in File S6.

**Table 5:** Closest relatives to the Gambian poultry strains

| 7-gene ST | cgST HC100 sub-cluster | Study poultry host | Neighbour host | Neighbour's country of isolation | Allelic distance |
| --- | --- | --- | --- | --- | --- |
| ST9286 | NA | Guinea fowl | Chicken | Gambia (this study) | 945 |
| ST9285 | NA | Chicken | Guinea fowl | Gambia (this study) | 945 |
| ST10654 | NA | Guinea fowl | Unknown avian source | Kenya | 1324 |
| ST155 | 43137 | Chicken and Guinea fowl | Poultry | US | 32-34 |
| ST2772 | NA | Chicken | Human | Kenya | 41 |
| ST6186 | NA | Chicken | Livestock | US | 58 |
| ST540 | 10207 | Guinea fowl | Human | UK | 59 |
| ST58 | 25133 | Chicken | Unknown | Unknown | 59 |
| ST2461 | 93699 | Chicken | Human | Kenya | 64 |
| ST2165 | 12281 | Chicken | Food | Kenya | 66 |
| ST4392 | NA | Guinea fowl | Human | UK | 68 |
| ST602 | NA | Chicken | Livestock | US | 70 |
| ST540 | 70056 | Chicken | Food | UK | 72 |
| ST540 | 1320 | Guinea fowl | Poultry | US | 73 |
| ST942 | NA | Guinea fowl | Environment (tap water) | Australia | 76 |
| ST212 | NA | Guinea fowl | Seagull | Australia | 81 |
| ST5826 | NA | Chicken | Water | UK | 91 |
| ST1423 | 27957 | Chicken | Reptile | US | 96 |
| ST337 | 73054 | Chicken | Reptile | US | 96 |
| ST196 | NA | Guinea fowl | Human | Kenya | 102 |
| ST155 | 93719 | Chicken | Tanzania | Human | 106 |
| ST86 | NA | Guinea fowl | US | Livestock | 131 |
| ST155 | 73905 | Guinea fowl | Companion animal | US | 137 |
| ST542 | 93732 | Chicken | Poultry | US | 148 |
| ST746 | NA | Chicken | Poultry | US | 148 |
| ST295 | NA | Guinea fowl | Human | Mexico | 162 |
| ST48 | 93724 | Chicken | Unknown | UK | 163 |
| ST542 | 93697 | Chicken | Environment (soil/dust) | US | 194 |
| ST155 | 73903 | Guinea fowl | Nepal | Human | 195 |
| ST443 | 93721 | Guinea fowl | Unknown | Unknown | 224 |
| ST6025 | NA | Guinea fowl | Unknown | US | 245 |
| ST2614 | NA | Guinea fowl | Human | China | 284 |
| ST9284 | NA | Chicken | Environment (soil/dust) | North America | 293 |
| ST2067 | NA | Guinea fowl | Human | Gambia | 458 |

**Figure 5:**
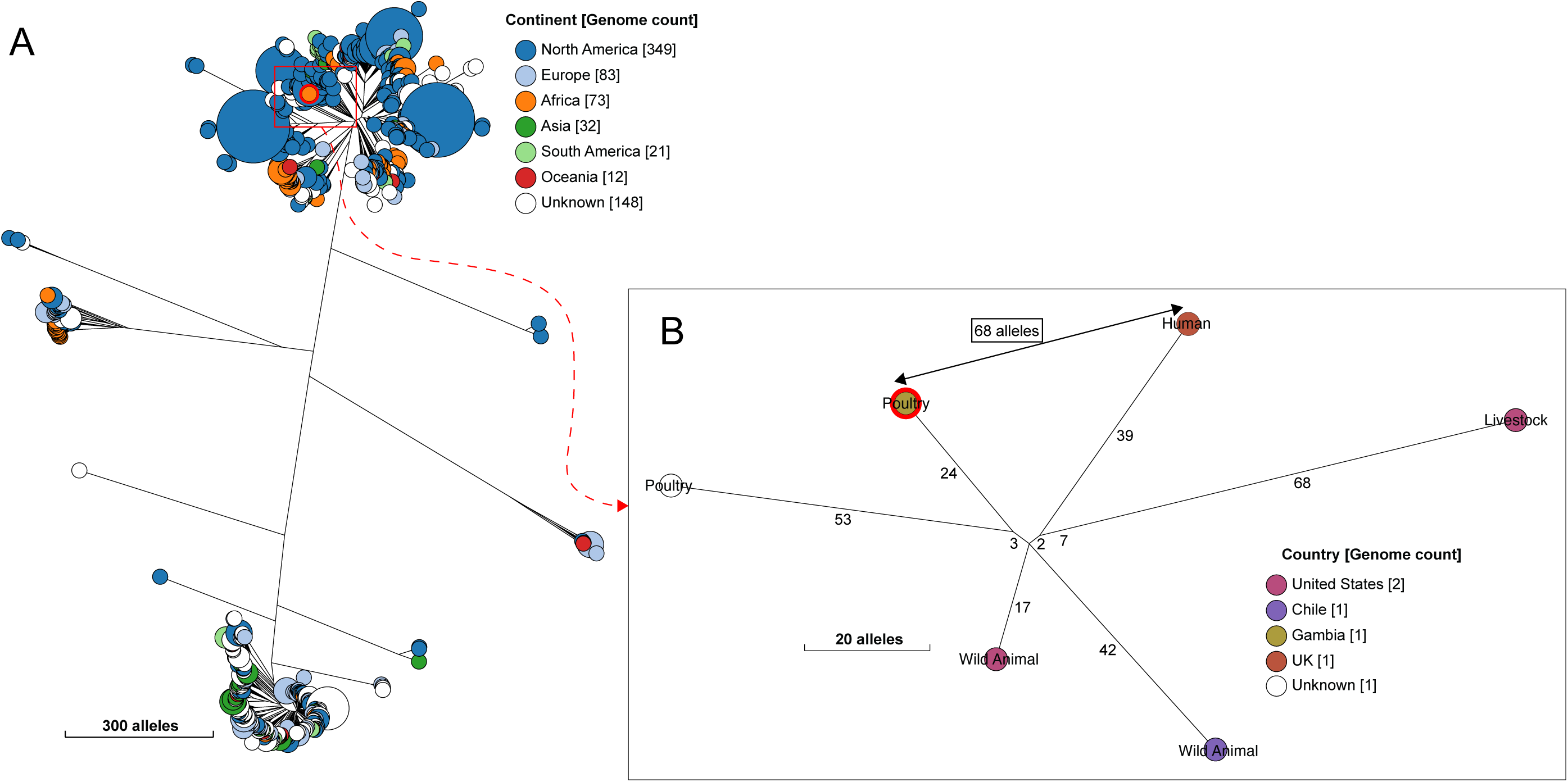
A NINJA neighbour-joining tree showing the phylogenetic relationship between the avian ST4392 (Achtman) strain from this study and all other publicly available genomes that cluster together at HC1100 level (cgST complex). The legend shows the continent of isolation of the isolates, with genome counts displayed in square brackets. Gambian poultry strains are highlighted in red. The study ST strain is separated from a human ST4392 isolate by 68 alleles, as shown in the subtree (B).

By contrast, three of the novel STs from this study (ST10654, ST9285, ST9286) were genetically distinct from anything else in the public domain. These belonged to unique HC1100 clusters in the cgMLST scheme and did not have any relatives in the seven-allele MLST scheme, even after allowing for two mismatches. Two of these (ST10654 from Guinea fowl 3 and ST9286 from Guinea fowl 4) now have complete genomic assemblies.

### The global prevalence of strains and AMR among avian *E. coli* isolates

Phylogenomic analyses of 4,846 poultry *E. coli* isolates from all over the world revealed that ST155 is common among poultry isolates from Africa and South America (Figures S1 and S2). In contrast, ST117 is prevalent among poultry isolates from Europe and North America (Figures S3-S4), with ST156 and ST254 being the most common *E. coli* STs found in poultry from Asia and Oceania respectively (Figures S5 and S6).

Our phylogenetic analyses revealed that ST155 strains from Africa were dispersed among other ST155 isolates from the rest of the world; however, the majority of ST155 strains from this study belonged to a tight genomic cluster, comprised of isolates from poultry and livestock from sub-Saharan Africa (separated by 38-39 alleles), except for a single isolate sourced from poultry in the US. In the cgMLST scheme, all the study ST155 isolates fell into four HC100 sub-clusters (100 alleles difference) (Figure S7). The largest sub-cluster (sub-cluster 1, HC100_43137) comprised ST155 isolates from this study and isolates from Uganda and Kenya; while sub-clusters 2 (HC100_73903), 3 (HC100_73905) and 4 (HC100_93719) occurred in the Gambia only—although distantly related to isolates from humans and a companion animal (Figure S8).

Antimicrobial resistance was high across the continents, with the highest prevalence of MDR in South America (100/131, 77%), followed by Asia (175/249, 70%), then Africa (392/591, 66%) (Table 4; File S8). Of note, the highest percentages of resistance globally were that of broad-spectrum beta-lactamases, while the lowest percentages of resistance were to colistin (File S6). Interestingly, the prevalence of colistin resistance was highest in Europe but did not occur in Oceania and North America.

## Discussion

Here, we have described genomic diversity of *E. coli* from backyard chickens and guinea fowl reared in households in rural Gambia, West Africa. Backyard poultry from this rural setting harbour a remarkably diverse population of *E. coli* strains that encode antimicrobial-resistance genes and virulence factors important for infections in humans. Furthermore, we provide evidence of sharing of strains (including MDR strains) from poultry to poultry and between poultry, livestock and humans, with potential implications for public health.

Our results reflect the rich diversity that exists within the *E. coli* population from backyard poultry. Although our sample size was small (19 birds), we recovered as many as 28 sequence types of *E. coli*, four of which have not been seen before—even though more than quarter of a million of *E. coli* strains had been sequence-typed by March 2020. Three of our novel STs differed by >945 alleles from their nearest relative—two of these now have complete assemblies. Also, some of the strains from this study were found in unique cgMLST HierCC clusters containing strains only from this study.

Our results confirm previous reports that phylogroups B1 and A are dominant phylogroups among *E. coli* isolates from both intensive and backyard poultry [54-57]. Hierarchical clustering analysis suggested that ST155 is common in African poultry. However, most of our ST155 strains belong to a unique cgMLST cluster containing closely related (38-39 alleles differences—and so presumably recently diverged) isolates from poultry and livestock from sub-Saharan Africa, suggesting that strains can be exchanged between livestock and poultry in this setting.

Rural backyard poultry can act as a source of transmission of infections to humans, due to the absence of biosecurity and daily contact with humans [58]. Indirect contact might occur through food or through contact with faeces, for example by children who are often left to play on the ground [59].

We observed a high prevalence of AMR genes among *E. coli* isolates sourced from African poultry. Similarly, high rates of genotypic MDR were detected among poultry *E. coli* isolates from the rest of the world, with ESBL (various types) being the most significant resistant gene detected. Poultry-associated ESBL genes have also been found among human clinical isolates [60]. Strikingly, most of our ST155 isolates encoded resistance to ≥3 classes of clinically relevant antibiotics, with the highest percentages to *bla-*TEM-1 beta-lactamase and tetracycline. This is worrying, as beta-lactamase-positive isolates are often resistant to several other classes of antibiotics [61, 62].

Our results are consistent with previous studies that reported ST155 isolates to be commonly associated with MDR [63, 64], but differ from other studies that have reported a low prevalence of AMR in backyard poultry. For example, in a study that compared the prevalence of ESBL genes in backyard poultry and commercial flocks from West Bengal, India, none of the 272 *E. coli* isolates from backyard birds harboured any ESBL gene [65], while 30% of commercial birds carried ESBL genes. The absence of resistance in that study was attributed to a lack of exposure to antimicrobials. Similarly, *E. coli* from organic poultry in Finland were reported to be highly susceptible to most of the antimicrobials studied and no ESBL resistance was detected [66].

Although tetracycline is commonly used in poultry farming for therapeutic purposes [67], resistance to this antibiotic is known to be prevalent in poultry, even in the absence of the administration of this antibiotic [68]. Our results also suggest that IncF plasmids may play a role in the dissemination of AMR in our study population.

Many sub-Saharan countries lack clear guidelines on the administration of antibiotics in agriculture, although an increasing trend in the veterinary use of antimicrobials has been documented [69]. The usage of antimicrobials in developing countries is likely to increase because of increasingly intensive farming practices [70]. Europe has banned the use of antimicrobials as growth promoters since 2006 [71] and the use of all essential antimicrobials for prophylaxis in animal production since 2011 [72]. However, AMR may be less well controlled in other parts of the world.

Although APEC strains span several phylogroups (A, B1, B2 and D) and serogroups [52], the majority of APEC strains encode virulence genes associated with intestinal or extra-intestinal disease in humans. These include adhesion factors, toxins, iron-acquisition genes, and genes associated with serum resistance—such as *fyu*A, *iuc*D, *iro*N, *iss, irp*2, *hly*F, *vat, kps*M and *omp*T. Although APEC isolates present different combinations of virulence factors, each retains the capability to cause colibacillosis [21, 73]. We did not detect haemolysin or serum survival genes in our study isolates; however, we recovered some of the known markers of intestinal and extraintestinal virulence in some study isolates—such the enteroaggregative *E. coli* heat-stable enterotoxin and the vacuolating autotransporter toxin (*vat, ast*A), invasion and evasion factors (*kps*M, *kps*D, *pla*) and adherence factors (*fim* and *pap* genes) that are associated with intestinal and extraintestinal infections in humans. Thus, these strains could cause disease in humans, should they gain access to the appropriate tissues.

Several birds were colonised with two or more STs and at least two phylotypes of *E. coli.* This level of diversity is probably a consequence of the frequent exposure of backyard poultry to the environment, livestock and humans. Co-colonisation of single hosts with multiple strains may facilitate the spread of AMR- and virulence-associated genes from resistant strains to other bacteria via both horizontal and vertical gene transfer [74]. A high co-colonisation rate of *E. coli* has been described in humans [75, 76] and in non-human primates [30]—involving pathogenic strains of *E. coli*. Recently, Li *et al* reported three to nine sequence types of colistin-resistant *E. coli* to co-exist within a single broiler chicken [28]. Here, we report co-colonisation with different lineages of *E. coli* in backyard chickens and guinea fowl. Unsurprisingly, co-colonising strains often had different AMR and virulence patterns.

An obvious limitation of our study is the small sample size. However, the inclusion of publicly-available sequences strengthens our analysis and inference of the population of *E. coli* in this setting. We also could not perform phenotypic susceptibility testing on all isolates. We acknowledge that a minor percentage of genotypic resistance predictions fail to correspond with phenotypic resistance [77].

Taken together, our results indicate a rich diversity of *E. coli* within backyard poultry from the Gambia, characterised by strains with a high prevalence of AMR and the potential to contribute to infections in humans. This, coupled with the potential for the exchange of strains between poultry and livestock within this setting, might have important implications for human health and warrants continued surveillance.

## Abbreviations

APEC: Avian Pathogenic *E. coli*;
ExPEC: Extraintestinal pathogenic *E. coli;*
ST: Sequence type;
AMR: Antimicrobial resistance;
MDR: Multiple-drug resistance;
MLST: Multi-locus sequence typing;
cgMLST: core-genome multi-locus sequence typing;
ARIBA: Antimicrobial resistance identification by assembly;;
VFDB: Virulence factors database;
SNP: single nucleotide polymorphism;
STGG: Skimmed milk tryptone glucose glycerol;
SDS: Sodium dodecyl-sulphate;
EDTA: Ethylenediaminetetraacetic acid;
ESBL: Expanded-spectrum beta-lactamase;
MIC: minimum inhibitory concentrations.

## Acknowledgements

We thank Dr Duto Fofana and Dr Ousman Ceesay of the Department of Livestock Services; Veterinary Services; Ministry of Agriculture, the Gambia for kindly providing access to their veterinary station at Abuko. We gratefully acknowledge the Research Support Office at the Medical Research Council Unit The Gambia at the London School of Hygiene and Tropical Medicine, in particular Mrs Elizabeth Stanley-Batchilly andsMs Isatou Cham, for project management support. We also thank Mr Steven Rudder and Mr David Baker for their assistance with the Illumina and Oxford nanopore sequencing.

## Author contributions

Conceptualization, MA, MP; data curation, MP, NFA; formal analysis, EFN; funding, MP and MA; sample collection, EFN, SJ, AS; laboratory experiments, EFN, supervision, BKA, AR, NFA, JO, MP, MA; manuscript preparation – original draft, EFN; review and editing, NT, NFA, MP; review of final manuscript, all authors.

## Supplementary Figures

**Figure S1.** A NINJA neighbour-joining tree of all publicly available *E. coli* poultry isolates from Africa, showing the prevalence of Achtman sequence types (STs). The dominant ST is highlighted with a red box. The legend displays the top 27 STs, with the respective genome counts displayed in square brackets.

**Figure S2.** A NINJA neighbour-joining tree of all publicly available *E. coli* poultry isolates from South America, depicting the prevalence of Achtman sequence types (STs). The most common ST found among *E. coli* isolates from this continent is ST155 (highlighted with a red box), similar to Africa (see Figure S1). The top 20 STs are displayed in the legend, with the respective genome counts displayed [in square brackets].

**Figure S3.** A NINJA neighbour-joining tree of all publicly available *E. coli* poultry isolates from Europe, depicting the prevalence of Achtman sequence types (STs). The top 20 STs are displayed in the legend, with the most common ST among poultry isolates from this continent (ST117) highlighted with a red box. The respective genome count per ST is also displayed.

**Figure S4.** A NINJA neighbour-joining tree of all publicly available *E. coli* poultry isolates from North America, showing the prevalence of Achtman sequence types (STs). The most common ST among poultry isolates from this continent is ST117 (highlighted with a red box). The legend displays the top 23 STs, with the respective genome counts displayed next to the STs.

**Figure S5.** A NINJA neighbour-joining tree of all publicly available *E. coli* poultry isolates from Asia, showing the prevalence of Achtman sequence types (STs). The most common ST among poultry isolates from this continent is ST156 (highlighted with a red box). The legend displays the top 25 STs, with the respective genome counts displayed next to the STs.

**Figure S6.** A NINJA neighbour-joining tree of all publicly available *E. coli* poultry isolates from Oceania, depicting the prevalence of Achtman sequence types (STs). The most common ST found among *E. coli* isolates from this continent is ST354 (highlighted with a red box). The first 18 STs are displayed in the legend, with the respective genome counts displayed.

**Figure S7.** A phylogenetic tree showing the global distribution of *E. coli* ST155 isolates. The study ST155 isolates are highlighted in red. Hierarchical clustering resolved four sub-clusters, encompassing the Gambian ST155 strains, displayed in red boxes. The legend displays the locations of the isolates, with the genome counts depicted in square brackets.

**Figure S8.** NINJA phylogenetic trees showing the sub-clusters for the study ST155 population within the cgMLST hierarchical clustering scheme. The largest sub-cluster (HC100_43137) (**A**) encompassed most of the study ST155 isolates (13/22, 59%), which were closely related to isolates from poultry and livestock in sub-Saharan Africa (separated by 38-39 alleles). Sub-clusters 2 (HC100_73903) (**B**), 3 (HC100_73905) (**C**) and 4 (HC100_93719) (**D**) were unique to the Gambia, although distantly related to isolates from humans and a companion animal. The red highlights indicate the study ST155 isolates. The locations of the isolates are displayed in the respective legends, with the genome counts indicated in square brackets.

**Figure S9.** NINJA phylogenetic trees showing the closest neighbours to avian ST86 isolates from this study. The nearest relatives occurred in livestock, depicted with the arrow. The legend indicates the location of isolation, with the genome count displayed in square brackets.

**Figure S10.** A NINJA phylogenetic tree showing the closest neighbours to avian ST6186 isolates from this study (A). The nearest relatives occurred in livestock from Kenya (B), separated by 58 alleles (depicted with the arrow). The branch lengths display the allelic distance between the genomes. The legend indicates the location of isolation, with the genome count displayed in square brackets.

**Figure S11.** A NINJA phylogenetic tree showing the closest neighbours to avian ST602 isolates from this study. The nearest relatives were isolated from livestock from the US (B), separated by 70 alleles (depicted with the arrow). The branch lengths display the allelic distance between the genomes. The legend indicates the location of isolation, with the genome count displayed in square brackets.

## Supplementary Files

**File S1.** Reference strains that were included in this study.

**File S2.** A summary of the sequencing statistics of the study isolates derived from this study.

**File S3. A.** A summary of the sequencing statistics of two novel sequence types derived from guinea fowl. **B.** Prophage types detected from long-read sequences using PHASTER (reference 50).

**File S4.** A pair-wise single nucleotide polymorphism matrix depicting the SNP distances between the study genomes calculated from the core genome alignment using snp-dists v0.6 (https://github.com/tseemann/snp-dists).

**File S5.** A summary of the predicted resistance genes for the study isolates, using ResFinder (Reference 24).

**File S6.** A list of the virulence factors detected among the study isolates using ARIBA VFDB (References 22 and 23) and their known functions.

**File S7.** Predicted plasmid-replicons from the study isolates.

**File S8.** The global prevalence of antimicrobial resistance genes predicted from whole-genome sequences of all publicly-available poultry *E. coli* isolates.

