## Supplementary material for "Genomic diversity of *Escherichia coli* isolates from backyard chickens and guinea fowl in the Gambia": File S1

**Supplementary File 1: Reference strains used in this study**

| Strain | Sequence Type | Phylogroup designation | | GenBank assembly accession |
| --- | --- | --- | --- | --- |
| K-12 strain MG1655 | ST10 | | A | NC_000913.3 |
| 536 | ST127 | | B2 | GCA_000013305.1 |
| UMN026 | ST597 | | D | GCA_000026325.2 |
| IAI39 | ST62 | | F | GCA_000026345.1 |
| O157:H7 str. EDL933 | ST11 | | E | GCA_000732965.1 |
| IAI1 | ST1128 | | B1 | GCA_000026265.1 |
| *Escherichia fergusonii* | ST5298 | | Outgroup | GCA_000026225.1 |
